# Representation of Protein Dynamics Disentangled by Time-structure-based Prior

**DOI:** 10.1101/2023.09.13.557264

**Authors:** Tsuyoshi Ishizone, Yasuhiro Matsunaga, Sotaro Fuchigami, Kazuyuki Nakamura

## Abstract

Representation learning (RL) is a universal technique for deriving low-dimensional disentangled representations from high-dimensional observations, aiding a multitude of downstream tasks. RL has been extensively applied to various data types, including images and natural language. Here, we analyze molecular dynamics (MD) simulation data of biomolecules in terms of RL to obtain disentangled representations related to their conformational transitions. Currently, state-of-the-art RL techniques, which are mainly motivated by the variational principle, try to capture slow motions in the representation (latent) space. Here, we propose two methods based on alternative perspective on the *disentanglement* in the representation space. The methods introduce a simple prior that imposes temporal constraints in the representation space, serving as a regularization term to facilitate capturing disentangled representations of dynamics. The introduction of this simple prior aids in characterizing the conformational transitions of proteins. Indeed, comparison with other methods via the analysis of MD simulation trajectories for alanine dipeptide and chignolin validates that the proposed methods construct Markov state models (MSMs) whose implied time scales are comparable to state-of-the-art methods. By coarse-graining MSMs, we further show the methods aid to detect physically important interactions for conformational transitions. Overall, our methods provide good representations of complex biomolecular dynamics for downstream tasks, allowing for better interpretations of conformational transitions.

## 1 Introduction

Molecular dynamics (MD) simulation is one of the most powerful approaches for investigating the conformational dynamics of biomolecules. ^1,2^ MD simulations can be used to investigate various dynamic phenomena of biomolecules, such as allosteric transitions,^3–8^molecular docking,^9–13^and structure formation.^14–18^Raw output data from MD simulations are trajectories, multivariate time-series data containing coordinates of atomic positions. The temporal information contained in trajectory data spans a broad range of time-scales, from atomic vibration in femtoseconds to protein folding in milliseconds. In the analysis of trajectory data, it is often challenging to reduce the high-dimensional data to a small number of collective variables (CVs), which summarize such complicated dynamics. Capturing good CVs is important for various downstream tasks including conformational clustering, free energy surface for analyzing the thermodynamics stability of states, Markov state modeling for analyzing kinetic behavior, and further simulations with enhanced sampling methods using the captured CVs.

For capturing CVs, dimensionality reduction techniques have been widely used in the studies of MD simulation analysis. Principal component analysis (PCA) captures the CVs with the largest variances by orthogonal transformations of the original coordinates. Relaxation mode analysis (RMA),^19,20^ time-structure-based independent component analysis (tICA),^21–25^ which are often applied for capturing CVs with the slowest relaxations by linear transformations. Advanced nonlinear reduction techniques include isomap ^26^and diffusion map.^27,28^ For details, the readers are referred to a recent excellent review on unsupervised learning of MD data by Glielmo et al.^29^

Generally, since the success of such downstream tasks is highly dependent on the choice of CVs, capturing “good” CVs is an important subject for analyzing trajectories. In terms of the field of machine learning, unsupervised learning techniques for obtaining good CVs or features with an emphasis on the performance of various downstream tasks have been studied in the context of representation learning (RL).^30–35^So far, RL has been developed and successfully applied to various types of data sets other than MD data. For image data, RL extracts informative representations for image classification, ^36,37^anomaly detection,^38,39^ and object detection.^40,41^ For language data, RL provides low-dimensional embedding for sentiment analysis,^42,43^named entity recognition,^44,45^ and language translation.^46^

The key point of RL is to learn features that *disentangle* many underlying explanatory factors (essential for downstream tasks, such as classification or prediction) hidden in the observed data as possible, discarding as little information about the data. ^30,31^ RL to obtain this kind of features include not only basic dimensionality reduction methods such as PCA but also dimensionality expansion methods such as sparse auto-encoder (SAE).^47^ In terms of this view, as shown by Schwantes et al.,^48–51^ tICA can be regarded as an optimal embedding of temporal variations in trajectory data by linear transformations. Recently, motivated by recent developments in deep learning technologies, a number of methods based on neural networks have been developed to disentangle the underlying temporal behavior hidden in trajectory data.^52–61^ An auto-encoder (AE) is a general nonlinear dimensionality reduction method based on neural networks that learn the process of reconstructing the original observed variables after encoding them into low-dimensional latent variables.^62^ Based on AE, various extensions of AE for MD trajectory data have been proposed. Time-lagged AE (TAE), which is an extension of AE to temporal domain, was developed to capture slow CVs by applying time-lagged reconstruction as the loss function in the AE framework.^52^ A variational AE (VAE) suppresses overfitting, which often occurrs in AE, by adding Gaussian noise to the latent variables.^63^ Time-lagged VAE (TVAE) learns encoding and time-lagged reconstruction process using VAE.^64^ Variational dynamics encoder (VDE) obtains slower CVs by penalizing TVAE’s loss function with a negative sample autocorrelation coefficient in the latent space.^53^ Gaussian-mixture VAE (GMVAE) simultaneously performs dimensionality reduction and clustering them into macrostates with a Gaussian mixture distribution.^60^ Variational approach for Markov processes nets (VAMPnets), state-free reversible VAMPnets (SRVs), and reversible deep MSM (revDMSM) learn macrostates based on a variational approach.^57,58,65^

Despite the development of these methods, the aspect of disentanglement in learning representations of dynamics has not been fully explored yet. Since the disentanglement is a key property for the success of downstream tasks, methods that can learn disentangled representations would be important for subsequent analyses. In order to explore this point, this study proposes two RL methods, time-structure-based variational autoencoder (tsVAE) and time-structure-based time-lagged variational autoencoder (tsTVAE). Both methods leverage a unique prior to disentangle complex biomolecular dynamics in the latent space. This prior imposes temporal constraints on the transitional motions in the latent space, expected to disentangle transitional motions on an event-by-event basis, not relying on averaged quantities like autocorrelations, which is a typical approach in other existing methods. Consequently, the proposed methods are capable of robustly learning disentangled representations from rel-atively short MD trajectory data, even when conformational transitions occur infrequently. Through the comparison with other existing methods on the MD simulation data of alanine-dipeptide and chignolin, we show that the proposed methods can extract disentangled CVs that capture conformational transitions well, and the learned representations (CVs) are further validated by the properties of constructed Markov state models.

This paper is organized as follows. Section 2 describes existing methods, then explain the details of the proposed methods (tsVAE, tsTVAE). Section 3 describes the protocols of MD simulations, and the properties of MSMs used to assess the qualities of RL methods. Section 4 show the results obtained by applying the existing and proposed methods to MD simulation data for alanine-dipeptide molecule and chignolin folding dynamics. Section 5 discusses the limits and the future directions of the proposed methods.

## 2 Theory

### 2.1 Notation

Let 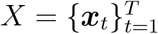 be the *d*_*x*_-dimensional *T* -length observed sequence and 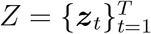 be the *d*_*z*_-dimensional *T* -length latent sequence. In our neural network model analysis, *X* is the feature-selected data from the series data obtained by MD simulation. For a vector sequence 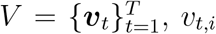, *v*_*t,i*_ represents the *i*-th element of the vector ***v***_t_. For a vector ***v****∈*ℝ^*d*^and a matrix *M∈*ℝ^*n×m*^, ***v***^*T*^*∈* ℝ^1*× d*^ and *M*^*T*^ *∈*ℝ^*m× n*^ represents the transpose of the vector and the matrix, respectively. For a vector ***v*** = (*v*_1_ ··· *v*_*d*_)^*T*^ *∈*ℝ^*d*^ and a natural number n *∈* ℕ, 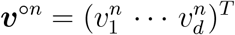 represents the element-wise power of n. For vectors ***v*** = (*v*_1_ *··· v*_*d*_)^*T*^ and ***u*** = (*u*_1_ *··· u*_*d*_)^*T*^ *∈*ℝ^*d*^, ***v ⊙ u*** = (*v*_1_*u*_1_ *··· v*_*d*_*u*_*d*_)^*T*^ represents the element-wise product. For a vector ***v*** = (*v*_1_ *··· v*_*d*_)^*T*^, 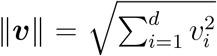 and 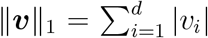 represent the L2 norm and the L1 norm, respectively. For a vector ***v****∈*ℝ^*d*^, diag(***v***) *∈* R^*d×d*^represents the diagonalized matrix. For a finite set S, |S| represents the number of elements of the set. For a random variable X *∈𝒳*, a function *f* : *𝒳 →* ℝ, and a distribution *p* : *𝒳 →* [0, ∞), 𝔼_*p*(*X*)_[*f*(*X*)] *∈* ℝ represents the expectation of *f*(*X*) regarding to the distribution *p*(*X*). For random variables *X ∈ 𝒳, Y ∈𝒴*, a function *f* : *𝒳 × 𝒴 →* ℝ, and a conditional distribution *p*(*X*|*Y* ), 𝔼 _*p*(*X*|*Y)*_[*f*(*X, Y* )] : *𝒳 →* ℝ represents the conditional expectation of *f*(*X, Y* ) regarding to the distribution *p*(*X*|*Y* ).

### 2.2 Auto-Encoder

Auto-encoder (AE) is a model for obtaining low-dimensional representation by reconstructing an observation from the reduced representation^66,67^(Figure 1(a)). The model embeds an observation ***x*** to the latent variable ***z*** = E *φ* (***x***) by the encoder 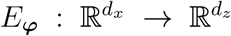 and reconstructs the observation 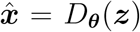 by the decoder 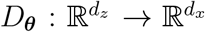 . The encoder E_***φ***_and the decoder *D*_θ_ are neural networks with the model parameters ***φ*** and ***θ***. The model parameters ***φ*** and ***θ*** are learned by minimizing the reconstruction loss

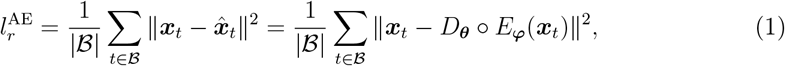

where *ℬ* is a minibatch whose elements are randomly sampled from *{*1, *···*,*T }*. Note that the AE does not use temporal information in the observations; thus it does not learn representations related to dynamic properties.

**Figure 1.**
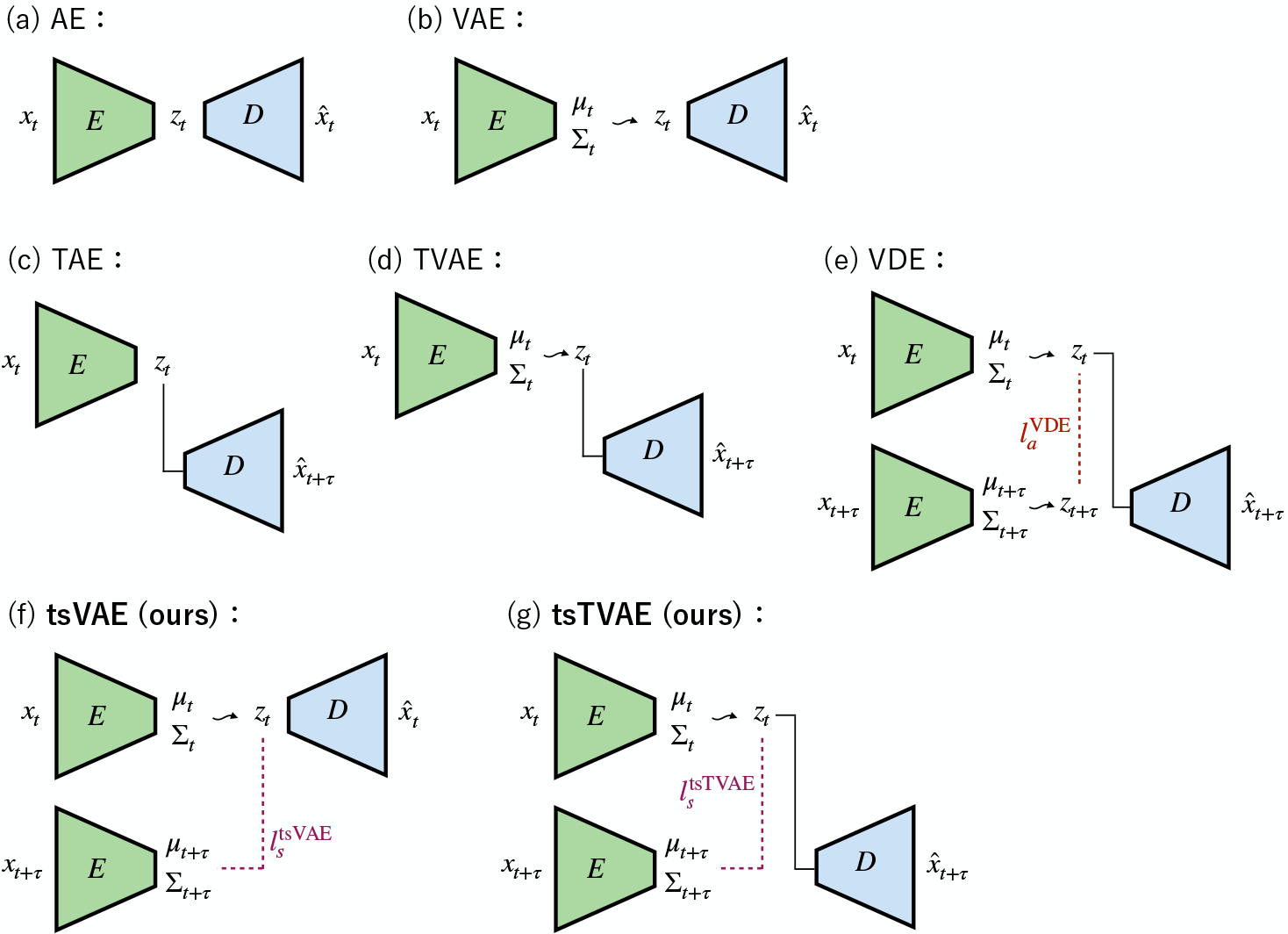
Neural network model structures of AE, VAE, TAE, TVAE, VDE, tsVAE, and tsTVAE. The green trapezoid E and the blue trapezoid D represent the encoder and the decoder, respectively. The red dashed line 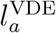, the purple dashed lines 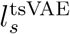, and 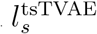represent the sample autocorrelation loss of VDE and the time-structure-based loss of tsVAE and tsTVAE, respectively. The black lines between a latent variable ***z***_t_ and the decoder D represent that the ***z***_t_ is an input of the decoder. The black wavy arrows represent sampling procedures from Gaussian distribution.

### 2.3 Variational Auto-Encoder

Variational auto-encoder (VAE) uses probabilistic models for AE’s encoder and decoder (Figure 1(b)). The model designs a variational posterior distribution *q*_*φ*_(***z***|***x***) and a generative distribution *p*_*θ*_ (***x***|***z***) by neural networks.^63^The parameters ***φ*** and ***φ*** are learned by variational inference,^68–70^ which is a method to maximize the evidence lower bound (ELBO) instead of intractable maximization of the log marginal likelihood. The ELBO is defined by

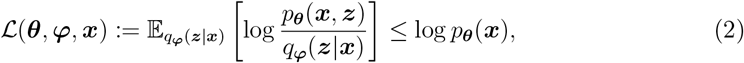

where *p*_*θ*_ (***x, z***) represents the joint generative distribution. The inequality in the equation (2) is easily proved by Jensen’s inequality.^71^The bound is represented by

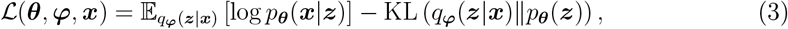

where *p*_***θ***_(***z***) is a prior distribution, and KL(*q*_*φ*_ (***z***|***x***)*‖p*_***θ***_(***z***)) represents the Kullback-Leibler (KL) divergence between the posterior q_φ_ (***z***|***x***) and the prior p_***θ***_ (***z***) distributions.^72–74^ As like the AE, the VAE does not learn the temporal information in the observations.

In a typical VAE, the prior *p*_***θ***_ (***z***) is chosen to be the standard isotropic Gaussian distribution *N*(***z***_*t*_; 0, *I*) where *N*(*·*; ***μ***, Σ) represents the normal distribution with mean vector ***μ*** and covariance Σ. This choice is computationally convenient (for calculating the KL divergence) and also encourages disentangling the representation in the latent space because the isotropic distribution promotes that latent variables become uncorrelated with each other. At the same time, the prior helps to reduce overfitting to limited sets of training data because the KL term in Eq. 3 works as a regularization term to constraint the capacity of information contained in the latent space.^75^

### 2.4 Time-lagged Neural Network Models

To learn the dynamic properties present in observations, the TAE extends the original AE by incorporating time-lagged information. The TAE^52^embeds an observation ***x***_*t*_ to the latent variable ***z***_*t*_ by the encoder network 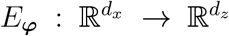 and reconstructs the time-lagged observation 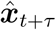 by the decoder network 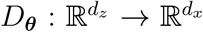 (Figure 1(c)). The model parameters ***φ*** and ***θ*** are learned by minimizing the reconstruction loss

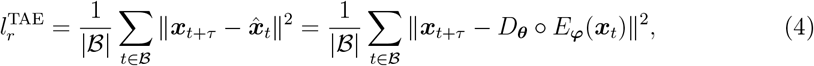

where ℬ*ℬ {*1, *···*,*T*−τ*}* and τ are a minibatch and a model lagtime, respectively.

The TVAE^64^ is an extension to TAE based on the architecture of the VAE. It introduces the probabilistic model to TAE and designs the variational posterior distribution, the generative distribution, and the prior distribution as follows (Figure 1(d)):

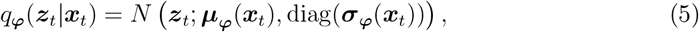

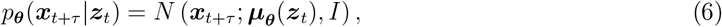

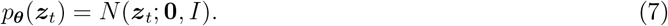

The model parameters ***φ*** and ***θ*** are learned by minimizing the following loss

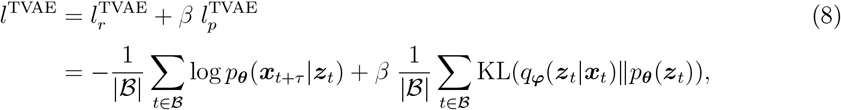

where 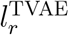 and 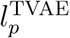 represent the reconstruction loss (negative log likelihood) and the prior regularized loss, respectively.

In the limit of a single linear hidden layer of the TAE, the tICA solution can be obtained.^52^ Also, the TVAE with regularized coefficient β = 0 reduces to the TAE with latent Gaussian perturbation. These suggest that the TAE and the TVAE are expected to be nonlinear extensions of tICA for learning of slow CVs hidden in high-dimensional observation. However, as discussed by Chen et al.,^59^ just minimizing the reconstruction loss does not ensure that the method capture the slowest CVs in nonlinear ways. Chen et al. theoretically show that, in general, the TAE learns a mixture of slow and maximum variance CVs instead of purely slow CVs.

The VDE^53^ is similar to the TVAE, but the problem of the TAE and TVAE is somewhat relaxed. Inspired by the variational approach to conformational dynamics,^76^ the VDE introduced the sample autocorrelation loss 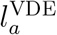 to the TVAE (Figure 1(e)). The VDE loss is defined by

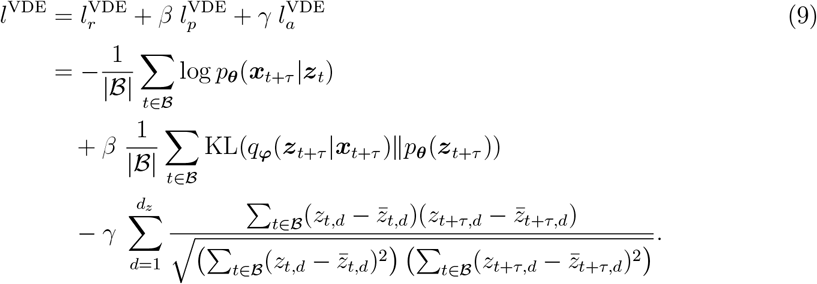

Here,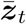 represents the mean of *{****z***_*t*_*}*_*t*∈*ℬ*_, and *ϒ* is an autocorrelation penalty coefficient. The sample autocorrelation loss 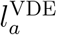 calculates the sum of the negative sample autocorrelation in each dimension; the smaller loss means higher sample autocorrelation in the latent space. This loss enhances obtaining slower CVs because higher autocorrelation corresponds to a slower transition in the latent space. While this loss term *ϒ* 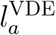 mainly depends on the coefficient *ϒ* and the minibatch size |*ℬ*|, the robustnesses are not fully discussed in the original paper.^53^

### 2.5 Time-structure-based Neural Network Models

Here, we describe our proposed RL methods, tsVAE and tsTVAE, designed to obtain disentangled representation of protein dynamics (Figure 1(f),(g)). The idea behind tsVAE and tsTVAE is based on the VAE is applying a prior *p*_***θ***_ (***z***) to encourage the disentanglement of its representation as done by the VAE. Whereas the notion of the VAE’s prior is limited to the disentanglement of static distributions, we extend the idea of the prior to temporal domain space. This prior works as temporal restraints, which encourages disentanglement of samples using temporal information. Specifically, the prior can be described as,

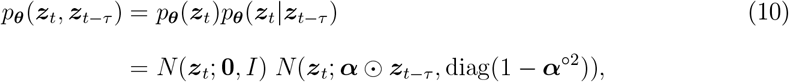

where 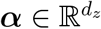 is a hyperparameter vector. The prior consists of the product of two normal distributions. The first normal distribution *N*(***z***_*t*_; 0, *I*) is the same as that used in the VAE and TVAE. As already mentioned above, it enhances disentanglement by isotropically distributing the latent variables. The second normal distribution *N*(***z***_*t*_; α ⊙ ***z***_*t*−τ_, diag(1 α^*;02*^)), which we refer to as *time-structure-based prior*, encourages the position of ***z***_*t*_ to be near time-proximal ***z***_*t*− τ_while maintaining a certain covariance around. All together, the second prior works to make time-proximal samples spatially close to each other while the first prior makes the other samples isotropically distributed or disentangled. Also, states that are far apart in time but visited many times in MD trajectories can be brought closer together by the reconstruction loss. Since the priors work as a regularization, it would be expected to avoid overfit to the transitional motions in fewer MD samples. This would make the tsVAE and tsTVAE robust for the analysis of a limited amount of MD data.

The model parameters ***φ*** and ***θ*** are learned by minimizing the negative bound

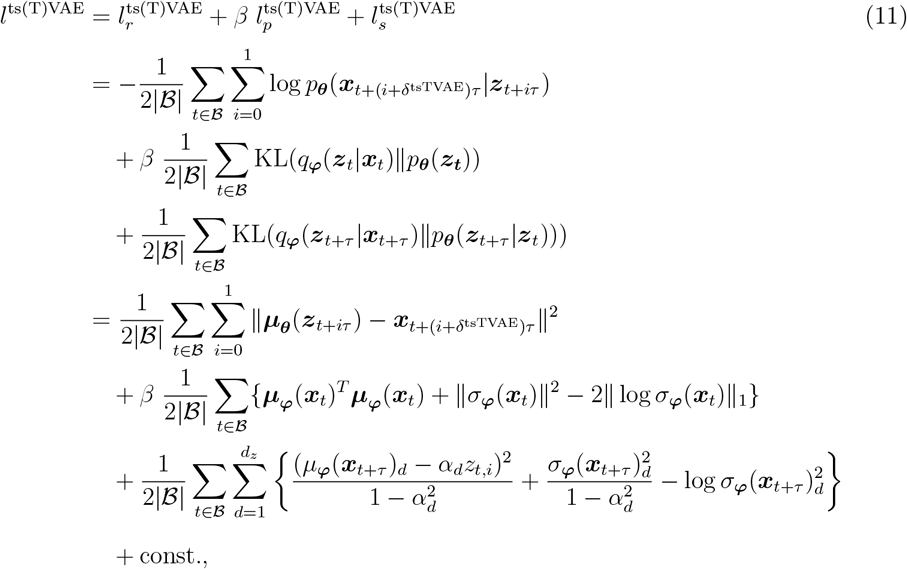

where δ^tsTVAE^ = 1 (tsTVAE); 0 (tsVAE) represent whether the model is the tsVAE or the tsTVAE. The third term (that came from the time-structure-based prior) in the equation works as a regularization on transitional motions during τ. The detailed training process for the tsVAE and the tsTVAE is shown in Algorithm 1.

#### Algorithm 1

Training of tsVAE and tsTVAE

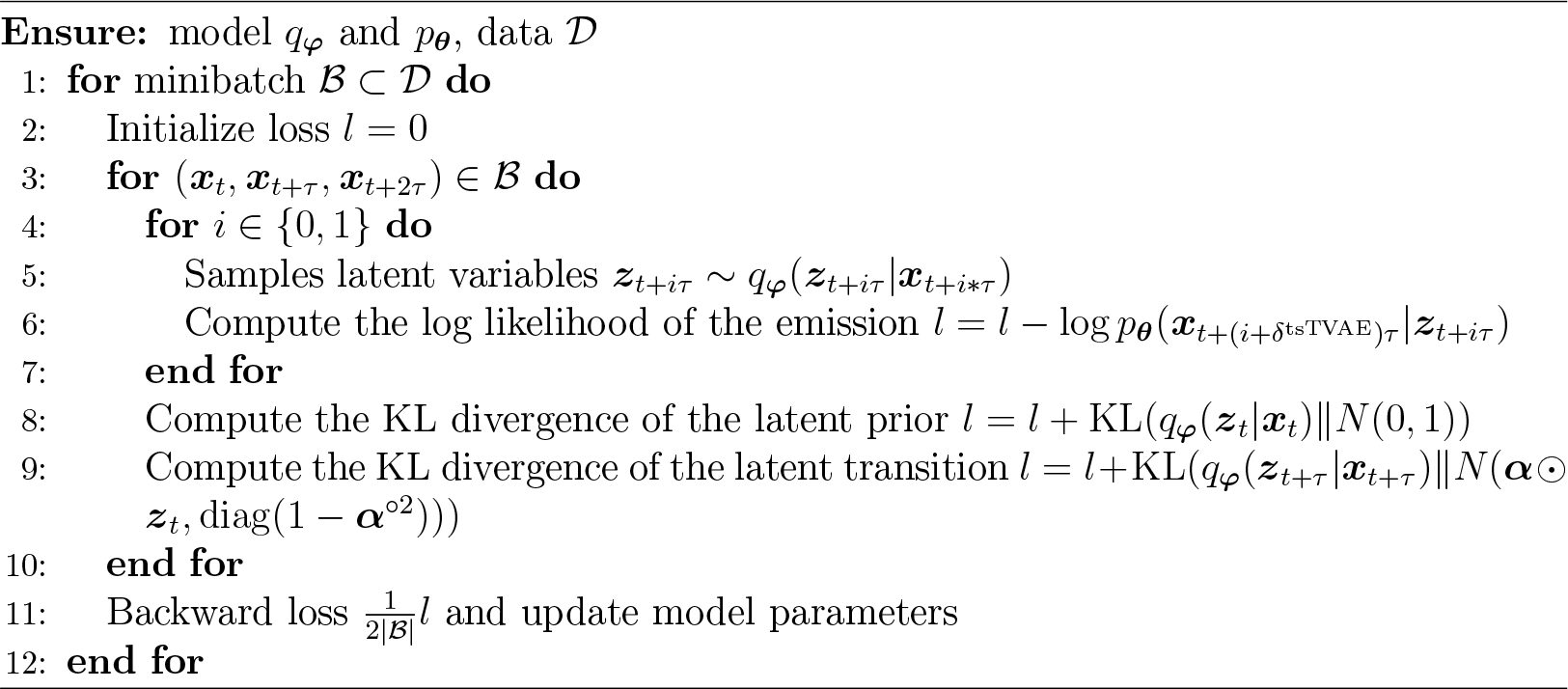

## 3 Methods

### 3.1 Molecular Dynamics Simulations

We compared RL methods through the analysis of MD trajectory data sets of alainine-dipeptide and chignolin. MD trajectory data of alanine-dipeptide^77,78^ was taken from a public data (https://markovmodel.github.io/mdshare/ALA2/#alanine-dipeptide) provided by the Computational Molecular Biology Group, Freie Universität Berlin. The total MD simulation length of the data is 250 ns performed using ACEMD.^77^ Amber ff99SB-ILDN force-field^79^ was used for proteins, and the TIP3P model^80^ was used for water molecules. All bonds involving hydrogen atoms were constrained.^81^ Electrostatic interactions were treated using the smooth particle mesh Ewald method.^82^ Temperature was controlled at 300 K by using Langevin dynamics.

MD trajectory data of chignolin was generated by ourselves through performing MD simulations. The total simulation length of 10 μs was performed with NAMD version 3.0 alpha (http://www.ks.uiuc.edu/Research/namd/).83 The input file for the MD simulation of chignolin was prepared by using the CHARMM-GUI web server.^84^ The initial structure was taken from PDB ID of 1UAO.^85^ The native structure was used as the initial state for the 200 ns simulation, and data were sampled every ten ps. CHARMM36m^86^ was used as the force field, and the TIP3P^80^ model was used for water molecules. The covalent bonds including hydrogen atoms were constrained.^81^ Electrostatic interactions were treated using the smooth particle mesh Ewald method.^82^ After equilibrating the system under NPT (300 K and 1 atm), a production run was conducted under NVT condition (300 K). The temperature was controlled by using Langevin dynamics. In the total 10 μs length simulation, five unfolding and four folding events were observed.

### 3.2 Markov State Model Analysis

We here used Markov state model (MSM)^87^ as a downstream task for the embedded representation. By constructing MSMs and assessing their properties, we compared the tsVAE and tsTVAE with PCA, tICA,^21–23^ TAE,^52^ TVAE,^64^ and VDE.^53^ The construction of MSM followed the standard procedure widely used in the community with the PyEMMA package:^88^

1. Feature selection. Aligned heavy-atom Cartesian coordinates were used for the alanine-dipeptide as features (*d*_*x*_ = 30). As preprocessing, the coordinates were z-scaled and transformed into triples 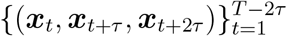 . For chignolin, the contact map vector was used as features. Specifically, the distance between alpha carbon atoms of non-adjacent residues was extracted from the MD data to obtain *d*_*x*_ = 28 data. The data was further transformed by exp(*-d*) and whitened to remove correlations between variables.
2. Representation learning. We embedded the selected features into the latent space by the PCA, tICA, TAE, TVAE, VDE, tsVAE, and tsTVAE.
3. Clustering. Embedded samples in the latent space by each method were clustered by the *k*-means method with *k* = 100, and discrete trajectories were obtained.
4. Construction of MSM. We constructed MSM by estimating a transition matrix by counting the number of transition events in the clustered discrete trajectories.
5. Coarse-graining or lumping MSM. The PCCA++^89^ algorithm was applied to compute macrostates decomposition of MSM states.

We compared the RL methods by assessing implied timescales (ITS) and eigenvectors of the transition matrix of MSM. The *i*-th ITS *t*_*i*_ is defined by

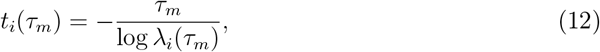

where τ_*m*_ is the MSM lagtime,, λ_*i*_ is the (*i* + 1)-th eigenvalue of the transition matrix. We further obtained macro-states by coarse-graining or lumping the constructed MSM with the PCCA++ algorithm.^89^We also defined ITS for this coarse-grained MSM, referring it as macro ITS.

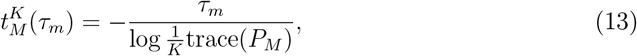

where *K* is the number of macrostates and *P*_*M*_ ∈ −^*k× k*^ is the macro transition probability matrix. We used the macro ITS to check whether slow time-scales (comparative to those of micro states) can be reproduced from macrostates. This is important assessment because if the macro ITS reproduce slow time-scales, it means that the latent space successfully disentangles transitional motions within broader regions of the space.

PCA and tICA were computed using the PyEMMA package,^88^ and the neural network models were implemented using PyTorch framework.^90^ The decoder network *D*_***φ***_ and the encoder network *E*_***θ***_ are composed of the number of hidden layers 2, respectively. Each layer has 50 units and the latent dimension is *d*_*z*_ = 2. We set the model lagtime τ = 50, the sample autocorrelation coefficient of VDE γ = 1, and the disentangled regularization coefficient β = 0. The neural network models were trained by using the Adam optimizer^91^ of a learning rate 10^3^, a batch size 256, and 100 epochs.

The hyperparameters ***α*** in the tsVAE and tsTVAE were determined by considering the decaying time τ_*d*_ of autocorrelation in the latent space. The decay time τ_*d*_ (*d* = 1, 2) and α_*d*_ (*d* = 1, 2) can be theoretically related by

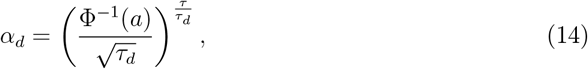

where a = 0.95 and Φ represent a significance level and the cumulative distribution function of the standard normal distribution, respectively. This relation means that the autocorrelation of lag time τ_*d*_ enters the accepted region of the uncorrelated hypothesis. In this study, we chose τ_*d*_ = 10^5^ (*d* = 1, 2) sampling steps for both alanine-dipeptide and chignolin and computed α_*d*_ (*d* = 1, 2) using this relation.

## 4 Results

### 4.1 Alanine-dipeptide

Figure 2 shows the comparison of MSM properties obtained from the alanine-dipeptide MD data. Since it is known that the alanine-dipeptide is well characterized by the two backbone dihedral angles (ϕ,ψ), we here investigated whether embedded spaces can achieve close correspondence to reference dihedral space (ϕ, ψ) only from the Cartesian coordinates (features used in MSM analysis). Figure 2(b) compares the convergence of the first three ITSs as a function of the MSM lagtime τ_*m*_. The ITSs of MSM using TAE, TVAE, VDE, tsVAE, and tsTVAE converge to the time scales comparable to those of the MSM constructed in the reference space (ϕ, ψ ). This means that these moethod successfully extract slow dynamics mainly determined by the dihedral angles (ϕ, ψ ). On the other hand, while the first two ITSs of MSM of tICA converge to the value comparable those of the reference space (ϕ, ψ ), the third implied timescale is significantly lower. In the case of PCA, the three ITSs of MSM are always lower than those of the reference space.

**Figure 2.**
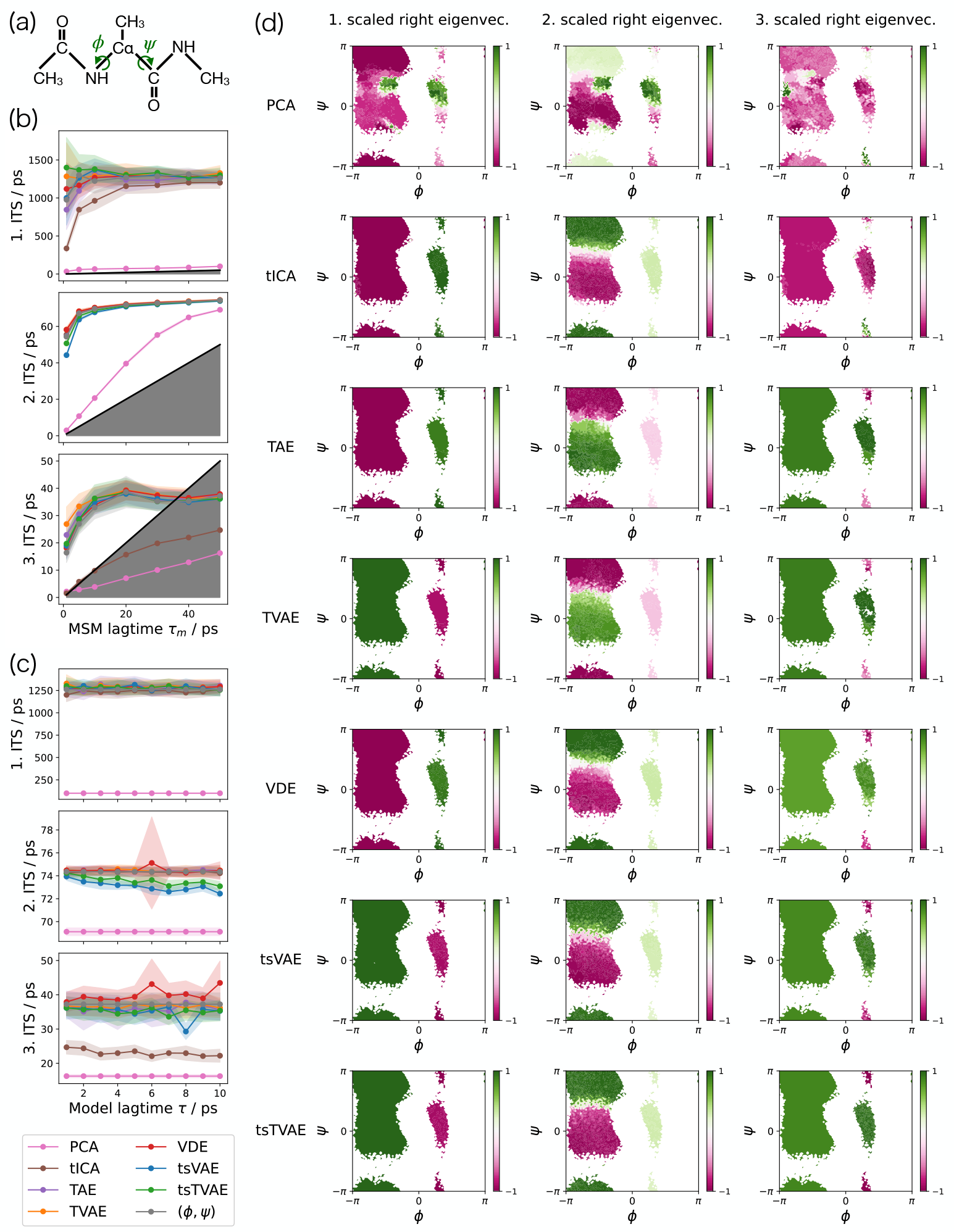
Comparison of Markov state models of alanine-dipeptide tajectory. (a) Structure of alanine-dipeptide. (b) First three implied timescales (ITSs) of MSMs constructed in the encoded space. (c) First two ITS at MSM lagtime 50 ps versus model lagtime τ. (d) The first three scaled right eigenvector maps of MSM with lagtime 50 ps in the reference space (ϕ, ψ).

Figure 2(c) shows the first three ITSs as a function of the model lagtime τ (while the MSM lagtime is fixed at 50 ps). The ITSs of TAE, TVAE, and VDE are robust to the choice of τ in [1 ps, 10 ps]. The tsVAE and tsTVAE have slight variations similar to those methods, while the second and the third ITS decrease with increasing τ . This is because the tsVAE and tsTVAE have a bias of reducing features on fast time-scales. It can be avoided by properly choosing the decay time τ_*d*_ and its corresponding hyperparameter ***α*** of these methods.

Next, we investigated whether the modes corresponding to the slowest three ITSs are well captured from the input Cartesian coordinates. Figure 2(d) shows the first three scaled right eigenvector maps in the reference space (ϕ, ψ ). The refereucen MSM shows that the first three slowest dynamics of the alanine-dipeptide are the ϕ-rotation, the -rotation with Gauche-negative ϕ, the ψ -rotation with Gauche-positive ϕ (more precisely, this shows the transition to left-handed helix region in the ABEGO-type Ramachandran plot), respectively. The three right eigenvectors of TAE, TVAE, VDE, tsVAE, and tsTVAE are consistent with those of the reference. On the other hand, PCA and tICA fail to capture the correspondent vectors, suggesting the superiority of non-linear transformations over these linear ones in obtaining a mapping from Cartesian coordinates to dihedral angles. Overall, these result confirms that the tsVAE and tsTVAE have comparable performance with the other state-of-the-art nonlinear methods in this simple system.

### 4.2 Chignolin

We evaluated the RL methods by analyzing the properties of constructed MSMs of chignolin’s folding/unfolding dynamics. Figure 3(a) shows representative structures of folded and unfolded states. The MD data for chignolin exhibits a more complex behavior than that of the alanine-dipeptide and is comparatively short, encompassing only five unfolding and four folding events within 10 μs length simulation. As such, this presents a more stringent comparison of the RL methods under consideration.

**Figure 3.**
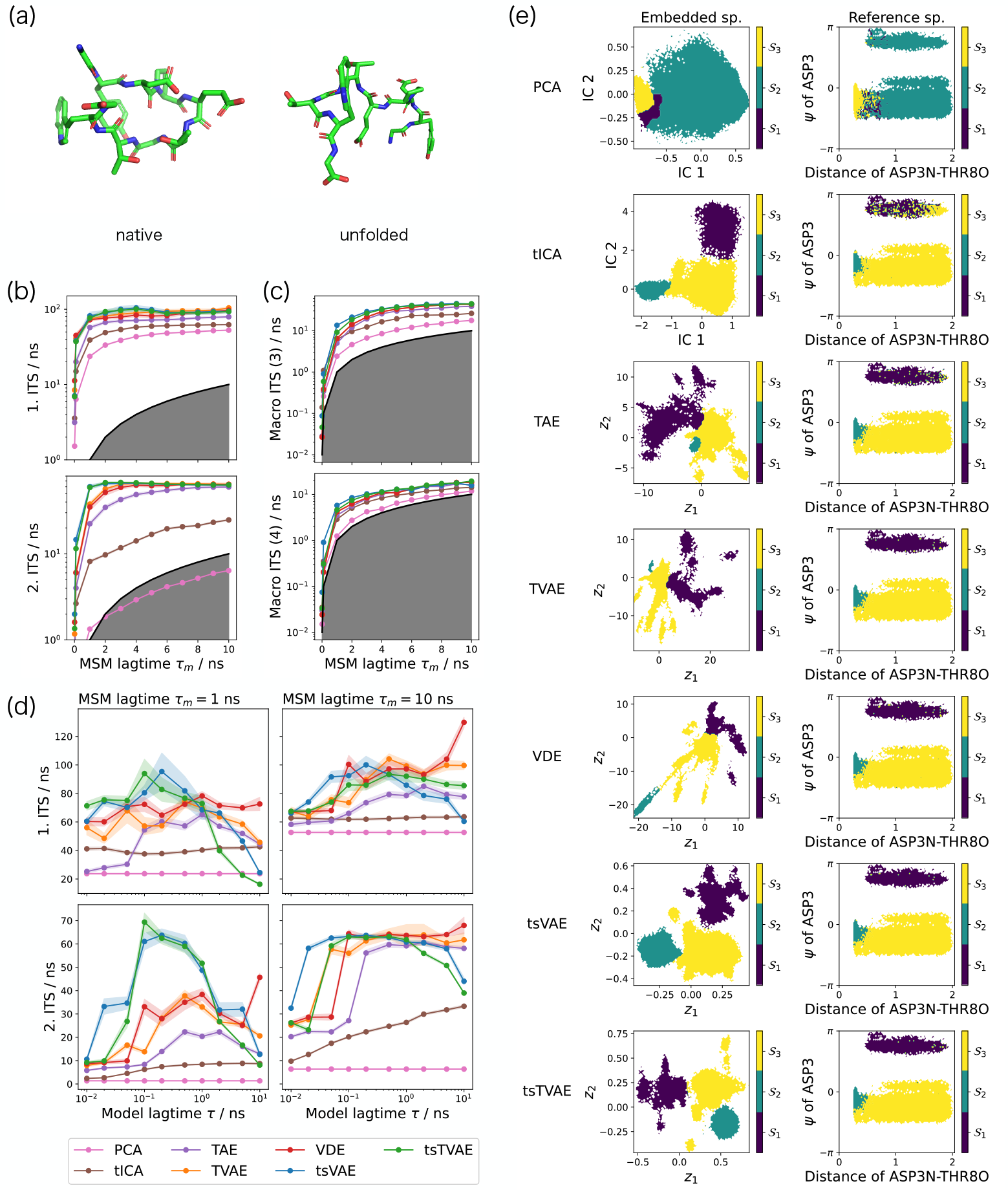
Comparison of Markov state models of chignolin folding and unfolding trajectory. (a) Representative structure of native and unfolded states. (b) First two implied timescales (ITSs) of MSMs constructed in the encoded space. (c) Macro ITSs of MSM. (d) First two ITSs at MSM lagtime 1 ns and 10 ns versus model lagtime τ . (e) Estimated macro-states of MSM with lagtime 1 ns in the embedded and the reference space.

Figure 3(b) depicts the first two ITSs of the MSMs constructed in the latent space. The figure shows that the ITSs of TAE, TVAE, VDE, tsVAE, and tsTVAE converge to slow timescales comparable with each other. In contrast, the 2nd ITSs of PCA and tICA do not converge to these time-scales. This indicates that nonlinear transformations are crucial for capturing the conformational transitions of chignolin. The figure also demonstrates that the 2nd ITSs of tsVAE and tsTVAE converge more rapidly than the other methods. This suggests that tsVAE and tsTVAE can robustly capture transitional motions, even when MD data is not long enough as like this case. This rapid convergence could be explained by the inductive bias due to the time-structure-based prior that samples closer in time will also be closer in the latent space.

We evaluated the robustness of each model against the choice of the model lagtime τ by varying τ (while fixing the MSM lagtime τ_*m*_ at 1 ns and 10 ns.) As shown in Figure 3(d), the ITSs of PCA, tICA, and TAE are significantly lower than those of other methods. In contrast, the ITSs of TVAE and VDE (with MSM lagtime τ_*m*_ = 10 ns) remain robustly high across a wide range from τ = 10^1^ ns to τ = 10^3^ ns, suggesting the impact of the sampling from the inference distribution on reducing the fast time-scales without overfit to the data. Additionally, the figure demonstrates that the ITSs of tsVAE and tsTVAE, especially when the MSM lagtime τ_*m*_ = 1 ns, are significantly higher than those of other methods in the range from τ = 10 ^−1^ ns to τ = 1 ns, suggesting the impact of the time-structure-based prior. Indeed, the ITSs tend to become high for relatively small τ because small τ increase the impact of the time-structure-based prior, which embeds samples close in the time to close position in the latent space.

The degree of disentanglement across broader regions of the latent space was explored through the properties of coarse-grained macrostates. We evaluated whether slow time-scales (comparative to those of micro states) can be reproduced from these macrostates of coarse-grained or lumped MSMs. Figure 3(c) demonstrates that the macro ITSs of both tsVAE and tsTVAE converge more rapidly to slower time-scales compared to the other methods (although the time-scales are lower than those of the micro states). This suggests that, compared to the other methods, the tsVAE and tsTVAE disentangle transitional motions within broader regions of the latent space.

The disentanglement in the macrostates is visualized in Figure 3(e). In the figure, samples in the latent space are colored according to three macrostates. Since the native structure of chignolin is well characterized by hydrogen bonds between the backbone amide group of ASP3 and the carbonyl group of THR8 (ASP3N-THR8O), we used this distance as a reference feature. In addition, we used the dihedral angle ψ of ASP as a feature because this angle is found to be coupled to the folding and unfolding transition in our MD data. In the figure, the macrostates of PCA and tICA are considered to be mixed (i.e., entangled) since the states are overlapped in the reference feature space (the right column in Figure 3(e)). On the other hand, the transitions among macrostates of TAE, TVAE, VDE, tsVAE, and tsTVAE are well correlated with the formation of contact between ASP3 and THR8 and the dihedral angle ψ of ASP. This concludes that these RL methods successfully capture important physical interactions upon the folding and unfolding transitions. Upon closer examination of the figure, the figure also shows that the tsVAE looks clearly disentangling the folding and unfolding transitions without mixing them. Indeed, *z*_1_ and *z*_2_ in the tsVAE have better correspondence with the ψ -rotation compared to the other methods. This could be explained by the property of the tsVAE and tsTVAE; the time-structure-based prior promotes time-local autocorrelation for each axis while the representations of each axis were disentangled to improve reconstruction accuracy.

### 4.3 Robustness against the choice of hyperparameters

The tsVAE and tsTVAE have the hyperparameters ***α*** in their time-structure-based prior which the user should give in advance. The robustness of results against the choice of the parameters ***α*** is important for the practical use of the methods. To evaluate the robustness, we conducted additional MSM analysis for alanine-dipeptide and chignolin trajectories. Figure 4 shows ITSs of MSMs constructed in the latent space with different ***α***. As desribed in Eq. (14), ***α*** can be related to the decay time τ_*d*_ of autocorrelation in the latent space as decirbed in Eq. (14). The first three ITSs for alanine-dipeptide trajectories have similar values for τ_*d*_ = 10^3^ ps to τ_*d*_ = 10^7^ ps. The first two ITSs for chignolin trajectories also have qualitatively same values from τ_*d*_ = 10^2^ ns to τ_*d*_ = 10^4^ ns. Overall, these results suggest the robustness of tsVAE and tsTVAE against the choice of the hyperparameters ***α***.

**Figure 4.**
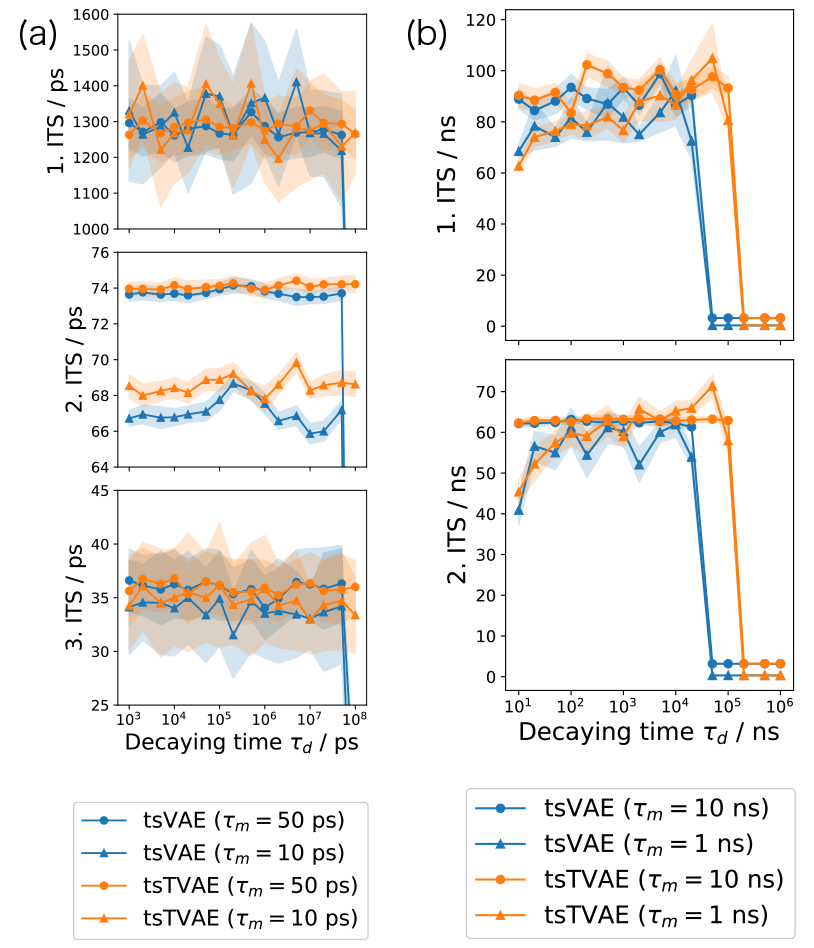
Implied timescales (ITS) of Markov state models with various choice of hyperparameters *α*. (a) The first three ITS at MSM lagtime 10 ps and 50 ps versus decaying time τ_d_ for alanine-dipeptide trajectories. (b) The first two ITS at MSM lagtime 1 ns and 10 ns versus decaying time τ_d_ for chignolin trajectories.

Furthermore, to eliminate dependency on ***α***, we introduced a prior for ***α*** and estimated plausible values of ***α*** from the data in Supporting Information. Another important hyperparameter is the learning rate η, used in the Adam optimizer for neural networks. The ITSs of tsVAE and tsTVAE are robustly higher than the other methods in η *∈* [10^−4^, 10 ^−2^]. Detailed results are given in Supporting Information.

## 5 Discussion

In this study, we have proposed two RL methods, tsVAE and tsTVAE, based on a simple time-structure-based prior. Through the comparison of MSM properties constructed from alanine-dipeptide’s MD trajectories, we have shown that the proposed methods can capture slow dynamics comparable to state-of-the-art methods. Furthermore, as shown by the macrostate analysis of coarse-grained MSMs for chignolin’s MD trajectories, the tsVAE or tsTVAE can learn disentangled representations that are well correlated with physically important interactions. This suggests that the proposed method can obtain widely disentangled representations leading to high interpretability. In the Supporting Information, we show a systematic analysis that can detect physically important interactions for the transitions between macrostates by analyzing the contributions of physical coordinates to the distributions of macro-states.

Although the tsVAE and tsTVAE methods are promising as alternatives for the current state-of-the-art methods, we would like to discuss limitations of these methods. The time-structure-based prior makes time-proximal samples spatially close to each other. Sometimes this effect introduces a bias into the dynamics in the latent space, leading to incorrect results in downstream tasks, including MSM analysis. For example, if the MD simulation is long enough to repeatedly visit the same metastable state in its simulation, but the visiting time is far apart from each other and missed to capture with the reconstruction loss, a bias is introduced by the time-structured-based prior by capturing this state as a different state in the latent space. To avoid this bias, we plan to develop an algorithm that iteratively clusters and learns embedding, such as SwAV.^92^

There would be two major future directions in this study. The first direction is to apply the methods for enhanced sampling of MD simulation. For example, in the Reweighted Autoencoded Variational Bayes for Enhanced Sampling (RAVE),^93^ and Deep-TICA,^94^ RL methods have been successfully combined with enhanced sampling techniques for biomolecular conformational sampling. Combination with the development version of OpenMM^95^ or PLUMED-LibTorch^96^ interface would be suitable to apply the biased forces from the latent variables of the tsVAE or tsTVAE as CVs for enhanced sampling.

The second direction is to extend the tsVAE and tsTVAE in the context of contrastive learning. Contrastive learning learns the general features of a dataset without labels by optimizing the embedding model to determine whether data points are similar or dissimilar. Since the time-structured-based prior of the proposed methods brings the latent variable ***z***_*t*_ and the time-proximal variable ***z***_*t*+τ_ closer together, the tsVAE and tsTVAE can be categorized to time contrastive learning.^97^ Thus, introducing the state-of-the-art techniques of contrastive learning in our context will further improve the representation of protein dynamics.

## Acknowledgement

This work was partly supported by JST CREST (Grant number: JPMJCR1762 to YM, and SF), and MEXT as “Program for Promoting Researches on the Supercomputer Fugaku” (Development and application of large-scale simulation-based inferences for biomolecules JPMXP1020230119 to TI, YM, and KN).

## Data Availability Statement

All of the code used in this study are publicly available at our GitHub repository https://github.com/ZoneTsuyoshi/time-structured-vae. The trajectory of chignolin folding and folding MD simulation is available from the corresponding author upon reasonable request.

## Supporting Information Available

The Supporting Information is available free of charge at XXX.

Additional analyses and comparisons of the representation learning methods (PDF)

## References

(1) Warshel, A.; Levitt, M. Theoretical studies of enzymic reactions – dielectric, electrostatic and steric stabilization of carbonium-ion in reaction of lysozyme. J. Mol. Biol. 1976, 103, 227–249.

(2) McCammon, J.; Gelin, B.; Karplus, M. Dynamics of folded proteins. Nature 1977, 267, 585–590.

(3) Monod, J.; Changeux, J.; Jacob, F. Allosteric proteins and cellular control systems. J. Mol. Biol. 1963, 6, 306–329.

(4) Monod, J.; Wyman, J.; Changeux, J. On nature of allosteric transitions – a plausible model. J. Mol. Biol. 1965, 12, 88–118.

(5) Koshland, D.; Nemethy, G.; Filmer, D. Comparison of experimental binding data and theoretical models in proteins containing subunits. Biochemistry 1966, 5, 365–385.

(6) Orellana, L.; Orozco, M. Understanding protein dynamics with coarsegrained models: from structures to disease. The FEBS Journal 2012, 279, 528–528.

(7) Sfriso, P.; Emperador, A.; Orellana, L.; Hospital, A.; Lluis, G. J.; Orozco, M. Finding conformational transition pathways from discrete molecular dynamics simulations. J. Chem. Theory Compute. 2012, 8, 4707–4718.

(8) Sfriso, P.; Hospital, A.; Emperador, A.; Orozco, M. Exploration of conformational transition pathways from coarse-grained simulations. Bioinformatics 2013, 29, 1980–1986.

(9) Maria Novoa, E.; Ribas de Pouplana, L.; Barril, X.; Orozco, M. Ensemble docking from homology models. J. Chem. Theory Compute. 2010, 6, 2547–2557.

(10) Fukunishi, Y. Structural ensemble in computational drug screening. Expert Opin. Drug Metab. Toxicol. 2010, 6, 835–849.

(11) Hawkins, P.; Nicholls, A. Conformer generation with OMEGA: learning from the data set and the analysis of failures. J. Chem. Inf. Model. 2012, 52, 2919–2936.

(12) Ishikawa, Y. A script for automated 3-dimentional structure generation and conformer search from 2-dimentional chemical drawing. Bioinformatics 2013, 9, 988–992.

(13) Klett, J.; Cortes-Cabrera, A.; Gil-Redondo, R.; Gago, F.; Morreale, A. ALFA: Automatic ligand flexibility assignment. J. Chem. Inf. Model. 2014, 54, 314–323.

(14) Ginalski, K.; Elofsson, A.; Fischer, D.; Rychlewski, L. 3D-jury: a simple approach to improve protein structure predictions. Bioinformatics 2003, 19, 1015–1018.

(15) Piana, S.; Lindorff-Larsen, K.; Shaw, D. Protein folding kinetics and thermodynamics from atomistic simulation. Proc. Natl. Acad. Sci. U.S.A. 2012, 109, 17845–17850.

(16) Raval, A.; Piana, S.; Eastwood, M.; Dror, R.; Shaw, D. Refinement of protein structure homology models via long, all-atom molecular dynamics simulations. Proteins 2012, 80, 2071–2079.

(17) Mirjalili, V.; Feig, M. Protein structure refinement through structure selection and averaging from molecular dynamics ensembles. J. Chem. Theory Compute. 2013, 9, 1294–1303.

(18) Dorn, M.; E Silva, M.; Buriol, L.; Lamb, L. Three-dimensional protein structure prediction: methods and computational strategies. Comput. Biol. Chem. 2014, 53, 251–276.

(19) Takano, H.; Miyashita, S. Relaxation Modes in Random Spin Systems. Journal of the Physical Society of Japan 1995, 64, 3688–3698.

(20) Mitsutake, A.; Takano, H. Relaxation mode analysis for molecular dynamics simulations of proteins. 10, 375–389.

(21) Naritomi, Y.; Fuchigami, S. Slow dynamics in protein fluctuations revealed by time-structure based independent component analysis: The case of domain motions. J. Chem. Phys. 2011, 134, 065101.

(22) Pérez-Hernández, G.; Paul, F.; Giorgino, T.; Fabritiis, G. D.; Noé, F. Identification of slow molecular order parameters for Markov model construction. J. Chem. Phys. 2013, 139, 015102.

(23) Schwantes, C. R.; Pande, V. S. Improvements in Markov State Model Construction Reveal Many Non-Native Interactions in the Folding of NTL9. J. Chem. Theory Compute. 2013, 9, 2000–2009.

(24) McGibbon, R. T.; Pande, V. S. Learning Kinetic Distance Metrics for Markov State Models of Protein Conformational Dynamics. J. Chem. Theory Compute. 2013, 9, 2900–2906.

(25) Sheong, F. K.; Silva, D.-A.; Meng, L.; Zhao, Y.; Huang, X. J. Automatic State Partitioning for Multibody Systems (APM): An Efficient Algorithm for Constructing Markov State Models To Elucidate Conformational Dynamics of Multibody Systems. J. Chem. Theory Compute. 2015, 11, 17–27.

(26) Das, P.; Moll, M.; Stamati, H.; Kavraki, L. E.; Clementi, C. Low-dimensional, freeenergy landscapes of protein-folding reactions by nonlinear dimensionality reduction. 103, 9885–9890, Publisher: Proceedings of the National Academy of Sciences.

(27) Singer, A.; Erban, R.; Kevrekidis, I. G.; Coifman, R. R. Detecting intrinsic slow variables in stochastic dynamical systems by anisotropic diffusion maps. 106, 16090–16095, Publisher: Proceedings of the National Academy of Sciences.

(28) Kim, S. B.; Dsilva, C. J.; Kevrekidis, I. G.; Debenedetti, P. G. Systematic characterization of protein folding pathways using diffusion maps: Application to Trp-cage miniprotein. 142, 085101.

(29) Glielmo, A.; Husic, B. E.; Rodriguez, A.; Clementi, C.; Noé, F.; Laio, A. Unsupervised Learning Methods for Molecular Simulation Data. 121, 9722–9758.

(30) Bengio, Y. Deep learning of representations: Looking forward. SLSP. 2013; pp 1–37.

(31) Bengio, Y.; Courville, A.; Vincent, P. Representation Learning: A Review and New Perspectives. IEEE TPAMI 2013, 35 .

(32) Siddharth, N.; Paige, B.; van de Meent, J.-W.; Desmaison, A.; Goodman, N. D.; Kohli, P.; Wood, F.; Torr, P. H. Learning Disentangled Representations with Semi-Supervised Deep Generative Models. NIPS. 2017.

(33) Chen, F.; Wang, Y.; Wang, B.; Kuo, C.-C. J. Graph Representation Learning: A Survey. APSIPA Transactions on Signal and Information Processing 2020, 9 .

(34) Wang, Y.; Hou, Y.; Che, W.; Liu, T. From static to dynamic word representations: a survey. IJMLC 2020, 11, 1611–1630.

(35) Tian, Y.; Sun, C.; Poole, B.; Krishnan, D.; Schmid, C.; Isola, P. What Makes for Good Views for Contrastive Learning? NeurIPS. 2020.

(36) Wang, P.; Han, K.; Wei, X.-S.; Zhang, L.; Wang, L. Contrastive Learning Based Hybrid Networks for Long-Tailed Image Classification. Proceedings of the IEEE/CVF Conference on Computer Vision and Pattern Recognition (CVPR). 2021; pp 943–952.

(37) Wang, X.; Yang, S.; Zhang, J.; Wang, M.; Zhang, J.; Yang, W.; Huang, J.; Han, X. Transformer-based unsupervised contrastive learning for histopathological image classification. Medical Image Analysis 2022, 81, 102559.

(38) Kopuklu, O.; Zheng, J.; Xu, H.; Rigoll, G. Driver Anomaly Detection: A Dataset and Contrastive Learning Approach. Proceedings of the IEEE/CVF Winter Conference on Applications of Computer Vision (WACV). 2021; pp 91–100.

(39) Chen, B.; Zhang, J.; Zhang, X.; Dong, Y.; Song, J.; Zhang, P.; Xu, K.; Kharlamov, E.; Tang, J. GCCAD: Graph Contrastive Learning for Anomaly Detection. IEEE Transactions on Knowledge and Data Engineering 2022, 1–14.

(40) Xie, E.; Ding, J.; Wang, W.; Zhan, X.; Xu, H.; Sun, P.; Li, Z.; Luo, P. DetCo: Unsupervised Contrastive Learning for Object Detection. Proceedings of the IEEE/CVF International Conference on Computer Vision (ICCV). 2021; pp 8392–8401.

(41) Sun, B.; Li, B.; Cai, S.; Yuan, Y.; Zhang, C. FSCE: Few-Shot Object Detection via Contrastive Proposal Encoding. Proceedings of the IEEE/CVF Conference on Computer Vision and Pattern Recognition (CVPR). 2021; pp 7352–7362.

(42) Xu, G.; Meng, Y.; Qiu, X.; Yu, Z.; Wu, X. Sentiment Analysis of Comment Texts Based on BiLSTM. IEEE Access 2019, 7, 51522–51532.

(43) Birjali, M.; Kasri, M.; Beni-Hssane, A. A comprehensive survey on sentiment analysis: Approaches, challenges and trends. Knowledge-Based Systems 2021, 226, 107134.

(44) Al-Moslmi, T.; Gallofré Ocaña, M.; L. Opdahl, A.; Veres, C. Named Entity Extraction for Knowledge Graphs: A Literature Overview. IEEE Access 2020, 8, 32862–32881.

(45) Li, J.; Sun, A.; Han, J.; Li, C. A Survey on Deep Learning for Named Entity Recognition. IEEE Transactions on Knowledge and Data Engineering 2022, 34, 50–70.

(46) Otter, D. W.; Medina, J. R.; Kalita, J. K. A Survey of the Usages of Deep Learning for Natural Language Processing. IEEE Transactions on Neural Networks and Learning Systems 2021, 32, 604–624.

(47) Le, Q. V.; Ranzato, M.; Monga, R.; Devin, M.; Chen, K.; Corrado, G. S.; Dean, J.; Ng, A. Y. Building High-Level Features Using Large Scale Unsupervised Learning. Proceedings of the 29th International Coference on International Conference on Machine Learning. Madison, WI, USA, 2012; p 507–514.

(48) Schwantes, C. R.; Pande, V. S. Modeling Molecular Kinetics with tICA and the Kernel Trick. J. Chem. Theory Compute. 2015, 11 .

(49) Harrigan, M. P.; Pande, V. S. Landmark Kernel tICA For Conformational Dynamics. bioRxiv.org e-Print archive 2017,

(50) Sultan, M. M.; Pande, V. S. tICA-metadynamics: accelerating metadynamics by using kinetically selected collective variables. J. Chem. Theory Compute. 2017, 13, 2440–2447.

(51) McGibbon, R. T.; Husic, B. E.; Pande, V. S. Identification of simple reaction coordinates from complex dynamics. J. Chem. Phys. 2017, 146, 044109.

(52) Wehmeyer, C.; Noé, F. Time-lagged autoencoders: Deep learning of slow collective variables for molecular kinetics. The Journal of Chemical Physics 2018, 148, 241703.

(53) Hernández, C. X.; Wayment-Steele, H. K.; Sultan, M. M.; Husic, B. E.; Pande, V. S. Variational encoding of complex dynamics. Physical Review E 2018, 97, 062412.

(54) Sultan, M. M.; Wayment-Steele, H. K.; Pande, V. S. Transferable neural networks for enhanced sampling of protein dynamics. arXiv preprint arXiv:1801.00636 2018,

(55) Bandyopadhyay, S.; Mondal, J. A deep autoencoder framework for discovery of metastable ensembles in biomacromolecules. J. Chem. Phys. 2021, 155, 114106.

(56) Doerr, S.; Ariz, I.; Harvey, M. J.; De Fabritiis, G. Dimensionality reduction methods for molecular simulations. arXiv preprint arXiv:1710.10629 2017,

(57) Mardt, A.; Pasquali, L.; Wu, H.; Noé, F. VAMPnets for deep learning of molecular kinetics. Nat. Commun. 2018, 8, 1–11.

(58) Chen, W.; Sidky, H.; Ferguson, A. L. Nonlinear discovery of slow molecular modes using state-free reversible VAMPnets. J. Chem. Phys. 2019, 150, 214114.

(59) Chen, W.; Sidky, H.; Ferguson, A. L. Capabilities and limitations of time-lagged autoencoders for slow mode discovery in dynamical systems. J. Chem. Phys. 2019, 151, 064123.

(60) Varolgüneş, Y. B.; Bereau, T.; Rudzinski, J. F. Interpretable embeddings from molecular simulations using Gaussian mixture variational autoencoders. Machine Learning: Science and Technology 2020, 1, 015012.

(61) Ward, M. D.; Zimmerman, M. I.; Swamidass, S.; Bowman, G. R. DiffNets: Selfsupervised deep learning to identify the mechanistic basis for biochemical differences between protein variants. bioRxiv.org e-Print archive 2020,

(62) Hinton, G. E.; Salakhutdinov, R. R. Reducing the dimensionality of data with neural networks. Science 2006, 313, 504–507.

(63) Kingma, D. P.; Welling, M. Auto-encoding Variational Bayes. International Conference on Learning Representations (ICLR). 2014.

(64) Hoffmann, M.; Scherer, M. K.; Hempel, T.; Mardt, A.; de Silva, B.; Husic, B. E.; Klus, S.; Wu, H.; Kutz, J. N.; Brunton, S.; Noé, F. Deeptime: a Python library for machine learning dynamical models from time series data. Machine Learning: Science and Technology 2021,

(65) Mardt, A.; Noé, F. Progress in deep Markov state modeling: Coarse graining and experimental data restraints. J. Chem. Phys. 2021, 155, 1–14.

(66) Rumelhart, D. E.; McClelland, J. L. Parallel Distributed Processing: Explorations in the Microstructure of Cognition: Foundations; 1987; pp 318–362.

(67) Yann LeCun, G. H., Yoshua Bengio Deep learning. Nature 2015, 521, 436–444.

(68) Beal, M. J. Variational Algorithms for Approximate Bayesian Inference; 2003.

(69) Jordan, M. I.; Ghahramani, Z.; Jaakkola, T. S.; Saul, L. K. An Introduction to Variational Methods for Graphical Models. Machine learning 1999, 37, 183–233.

(70) Zhang, C.; Butepage, J.; Kjellstrom, H.; Mandt, S. Advances in variational inference. 2017.

(71) Jensen, J. L. W. V. Sur les fonctions convexes et les inégalités entre les valeurs moyennes. Acta Mathematica 1906, 30, 175 –193.

(72) Kullback, S.; Leibler, R. A. On information and sufficiency. Annals of Mathematical Sciences 1951, 22, 79–86.

(73) Kullback, S. Information Theory and Statisticcs; John Wiley & Sons, 1959.

(74) Amari, S. Information Geometry and Its Applications. Applied Mathematical Sciences 2016, 194, 374.

(75) Higgins, I.; Matthey, L.; Pal, A.; Burgess, C.; Glorot, X.; Botvinick, M.; Mohamed, S.; Lerchner, A. beta-VAE: Learning Basic Visual Concepts with a Constrained Variational Framework. International Conference on Learning Representations (ICLR). 2017.

(76) Noé, F.; Nüske, F. A Variational Approach to Modeling Slow Processes in Stochastic Dynamical Systems. Multiscale Modeling & Simulation 2013, 11, 635–655.

(77) Harvey, M. J.; Giupponi, G.; Fabritiis§, G. D. ACEMD: Accelerating Biomolecular Dynamics in the Microsecond Time Scale. J. Chem. Theory Compute. 2009, 5, 1632–1639.

(78) Nüske, F.; Wu, H.; Prinz, J.-H.; Wehmeyer, C.; Clementi, C.; Noé, F. Markov state models from short non-equilibrium simulations—Analysis and correction of estimation bias. J. Chem. Phys. 2017, 146, 094104.

(79) Lindorff-Larsen, K.; Piana, S.; Palmo, K.; Maragakis, P.; Klepeis, J. L.; Dror, R. O.; Shaw, D. E. Improved side-chain torsion potentials for the Amber ff99SB protein force field. Proteins: Structure, Function, and Bioinformatics 2010, 78, 1950–1958.

(80) Jorgensen, W. L.; Chandrasekhar, J.; Madura, J. D.; Impey, R. W.; Klein, M. L. Comparison of simple potential functions for simulating liquid water. 79, 926–935, Publisher: American Institute of Physics.

(81) Ryckaert, J.-P.; Ciccotti, G.; Berendsen, H. J. Numerical integration of the cartesian equations of motion of a system with constraints: molecular dynamics of n-alkanes. Journal of Computational Physics 1977, 23, 327–341.

(82) Darden, T.; York, D.; Pedersen, L. Particle mesh Ewald: An Nlog(N) method for Ewald sums in large systems. The Journal of Chemical Physics 1993, 98, 10089–10092.

(83) Phillips, J. C.; Hardy, D. J.; Maia, J. D. C.; Stone, J. E.; Ribeiro, J. V.; Bernardi, R. C.; Buch, R.; Fiorin, G.; Hénin, J.; Jiang, W.; McGreevy, R.; Melo, M. C. R.; Radak, B. K.; Skeel, R. D.; Singharoy, A.; Wang, Y.; Roux, B.; Aksimentiev, A.; Luthey-Schulten, Z.; Kalé, L. V.; Schulten, K.; Chipot, C.; Tajkhorshid, E. Scalable molecular dynamics on CPU and GPU architectures with NAMD. 153, 044130.

(84) Jo, S.; Kim, T.; Iyer, V. G.; Im, W. CHARMM-GUI: A webbased graphical user interface for CHARMM. 29, 1859–1865, _eprint: https://onlinelibrary.wiley.com/doi/pdf/10.1002/jcc.20945.

(85) Honda, S.; Yamasaki, K.; Sawada, Y.; Morii, H. 10 residue folded peptide designed by segment statistics. Structure 2004, 12, 1507–1518.

(86) Huang, J.; Rauscher, S.; Nawrocki, G.; Ran, T.; Feig, M.; de Groot, B. L.; Grubmüller, H.; MacKerell, A. D. CHARMM36m: an improved force field for folded and intrinsically disordered proteins. 14, 71–73.

(87) Husic, B. E.; Pande, V. S. Markov State Models: From an Art to a Science. Journal of the American Chemical Society 2018, 140, 2386–2396, PMID: 29323881.

(88) Scherer, M. K.; Trendelkamp-Schroer, B.; Paul, F.; Pérez-Hernández, G.; Hoffmann, M.; Plattner, N.; Wehmeyer, C.; Prinz, J.-H.; Noé, F. PyEMMA 2: A Software Package for Estimation, Validation, and Analysis of Markov Models. Journal of Chemical Theory and Computation 2015, 11, 5525–5542.

(89) Röblitz, S.; Weber, M. Fuzzy spectral clustering by PCCA+: application to Markov state models and data classification. 7, 147–179.

(90) Paszke, A.; Gross, S.; Massa, F.; Lerer, A.; Bradbury, J.; Chanan, G.; Killeen, T.; Lin, Z.; Gimelshein, N.; Antiga, L.; Desmaison, A.; Kopf, A.; Yang, E.; DeVito, Z.; Raison, M.; Tejani, A.; Chilamkurthy, S.; Steiner, B.; Fang, L.; Bai, J.; Chintala, S. PyTorch: An Imperative Style, High-Performance Deep Learning Library. Advances in Neural Information Processing Systems 2019, 8024–8035.

(91) Kingma, D. P.; Ba, J. Adam: A Method for Stochastic Optimization. International Conference on Learning Representations (ICLR). 2014.

(92) Caron, M.; Misra, I.; Mairal, J.; Goyal, P.; Bojanowski, P.; Joulin, A. Unsupervised learning of visual features by contrasting cluster assignments. NeurIPS. 2020.

(93) Ribeiro, J. M. L.; Bravo, P.; Wang, Y.; Tiwary, P. Reweighted autoencoded variational Bayes for enhanced sampling (RAVE). The Journal of Chemical Physics 2018, 149, 072301.

(94) Bonati, L.; Piccini, G.; Parrinello, M. Deep learning the slow modes for rare events sampling. Proceedings of the National Academy of Sciences 2021, 118, e2113533118.

(95) Eastman, P.; Swails, J.; Chodera, J. D.; McGibbon, R. T.; Zhao, Y.; Beauchamp, K. A.; Wang, L.-P.; Simmonett, A. C.; Harrigan, M. P.; Stern, C. D.; Wiewiora, R. P.; Brooks, B. R.; Pande, V. S. OpenMM 7: Rapid development of high performance algorithms for molecular dynamics. 13, e1005659.

(96) Tribello, G. A.; Bonomi, M.; Branduardi, D.; Camilloni, C.; Bussi, G. PLUMED 2: New feathers for an old bird. Computer Physics Communications 2014, 185, 604–613.

(97) Hyvarinen, A.; Morioka, H. Unsupervised feature extraction by time-contrastive learning and nonlinear ICA. Neural Information Processing Systems (NIPS). 2016.

