## supplementary information for "Representation of Protein Dynamics Disentangled by Time-structure-based Prior"

### Supporint Information for Representation of Protein Dynamics Disentangled by Time-structure-based Prior

#### A Learning Model Autocorrelation

While we used the fixed hyperparameter $\alpha$ adjusted from the fixed decaying time $\tau_d$ in our main experiments, we propose another setting called learned mode. The learned mode of the

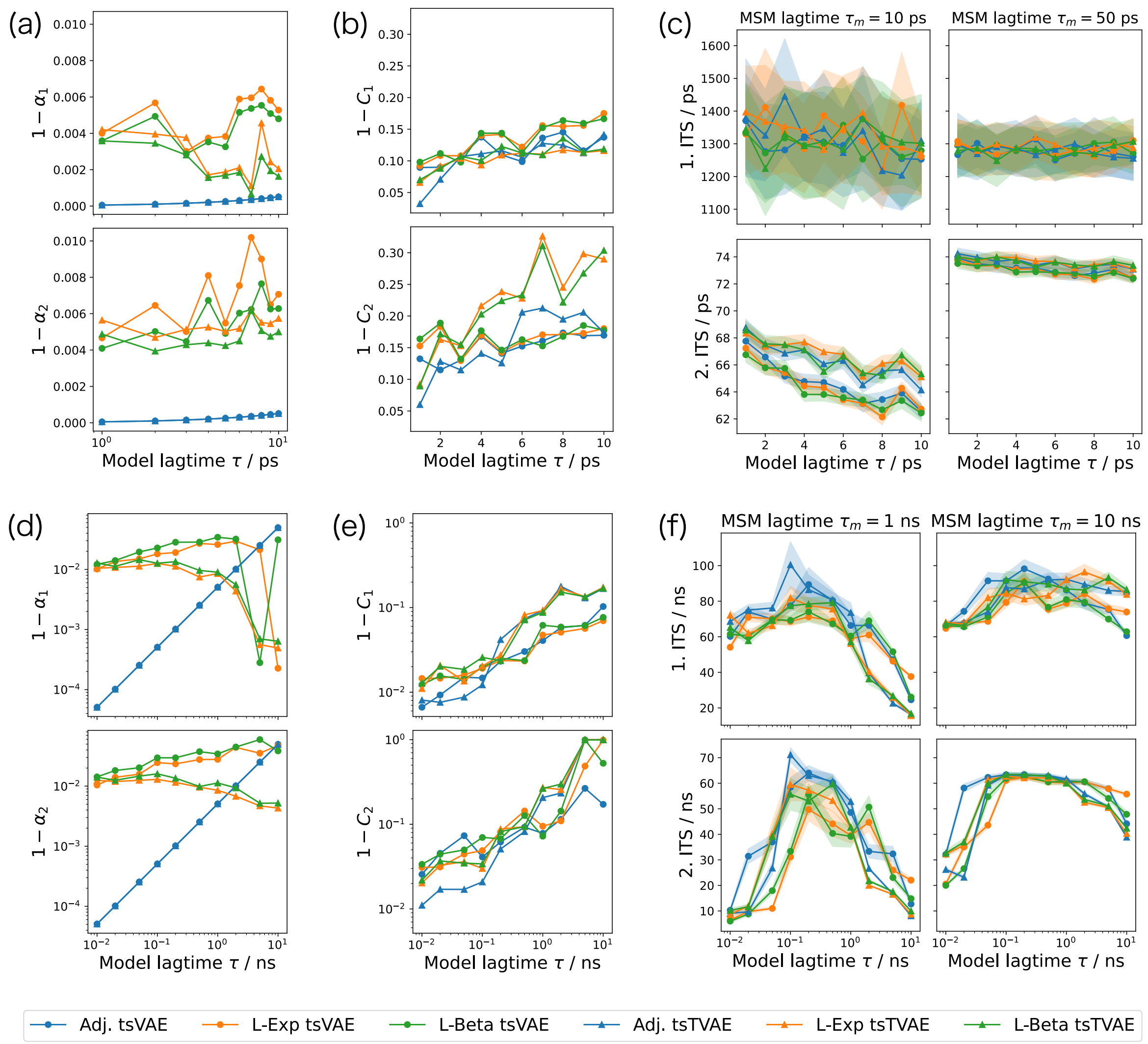


Figure 1: **Negative autocorrelations and implied timescales (ITS) of MSMs constructed in the latent space**. (a) The negative model autocorrelation $1-\alpha$ versus a model lagtime $\tau$ for alanine-dipeptide trajectories. (b) The negative sample autocorrelation $1-C$ versus a model lagtime $\tau$ for alanine-dipeptide trajectories (c) The first three ITS at MSM lagtime 50 ps versus a model lagtime $\tau$ for alanine-dipeptide trajectories. (d) The negative model autocorrelation $\boldsymbol{\epsilon} = 1-\boldsymbol{\alpha}$ versus a model lagtime $\tau$ for chignolin trajectories. (e) The negative sample autocorrelation $1-C$ versus a model lagtime $\tau$ for chignolin trajectories (f) The first three ITS at MSM lagtime 1 ns versus a model lagtime $\tau$ for chignolin trajectories.

model means the hyperparameter $\alpha$ is learned together with the neural network parameters. This mode assumes a prior distribution such as the Beta distribution[1,2] $B(3,1)$. The overall time-structured prior distribution is designed by

$$p_{\boldsymbol{\theta}}(Z,\boldsymbol{\alpha}) = p(\boldsymbol{\alpha})\prod_{t=1}^{\tau} p_{\boldsymbol{\theta}}(\boldsymbol{z}_t)\prod_{t=\tau+1}^{T} p_{\boldsymbol{\theta}}(\boldsymbol{z}_t|\boldsymbol{z}_{t-\tau},\boldsymbol{\alpha}) \quad (1)$$

$$= p(\boldsymbol{\alpha})\prod_{t=1}^{\tau} N(\boldsymbol{z}_t;\boldsymbol{0},I)\prod_{t=\tau+1}^{T} N(\boldsymbol{z}_t;\boldsymbol{\alpha}\odot\boldsymbol{z}_{t-\tau},\mathrm{diag}(1-\boldsymbol{\alpha}^{\circ 2})). \quad (2)$$

In the following experiment, we used the Beta distribution $B(3,1)$ and the pseudo-exponential distribution whose probability density function is defined by

$$p(\alpha) = e^{\alpha\log 3} - 1. \quad (3)$$

Figure 1 shows the negative autocorrelations and implied timescales (ITS) of MSMs constructed in the latent space. Figure 1 (a) and (d) show the negative model autocorrelation $\boldsymbol{\epsilon} = 1 - \boldsymbol{\alpha}$ for alanine-dipeptide and chignolin trajectories, respectively. For the adjusted mode, the parameters were computed by (14) of the main article. For the learned mode, the parameters were learned together with neural network parameters. The first three implied timescales of all modes for alanine-dipeptide trajectories are competitive (Figure 1 (c)). The first two implied timescales of the adjusted mode are relatively higher than those of the learned mode for chignolin trajectories (Figure 1 (f)). This means that under the known targetted actual timescale, the adjusted mode provided more suitable ITSs than the learned mode.

### B Robustness for Learning Rate

A learning rate is an important hyperparameter for optimizing neural network parameters. Figure 2 shows implied timescales of MSMs constructed in the latent space with each learning

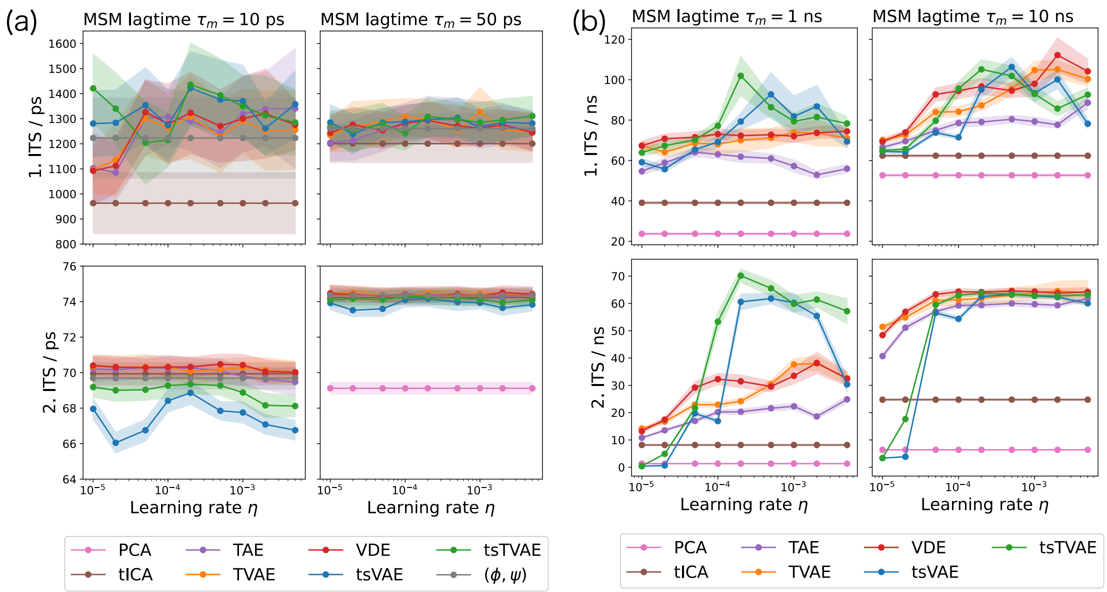


Figure 2: **Implied timescales (ITS) of MSMs constructed in the latent space**. (a) The first three ITS at MSM lagtime 10 ps and 50 ps versus learning rate $\eta$ for alanine-dipeptide trajectories. (b) The first two ITS at MSM lagtime 1 ns and 10 ns versus learning rate $\eta$ for chignolin trajectories.

rate. The first three implied timescales for alanine-dipeptide trajectories slightly vary for the learning rate (Figure 2(a)). The first two implied timescales of tsVAE and tsTVAE at MSM lagtime 1 ns for chignolin trajectories are higher than the other methods in $\eta \in [10^{-4}, 10^{-2}]$. This suggests that the proposed methods can obtain slow CVs more efficiently than the existing methods by exploring the learning rate scale.

### C How to Detect Key Variables from MSM Macro-states

Detecting key variables that contributed to slow dynamics is essential. To extract the variables, we used each variable's total variation distances between histograms of macro-states. The total variation distance between two probability distributions $p_1$ and $p_2$ is formulated by

$$\|p_1 - p_2\|_{\mathrm{TV}} = \sup_{A \in \mathcal{F}} |p_1(A) - p_2(A)|, \tag{4}$$

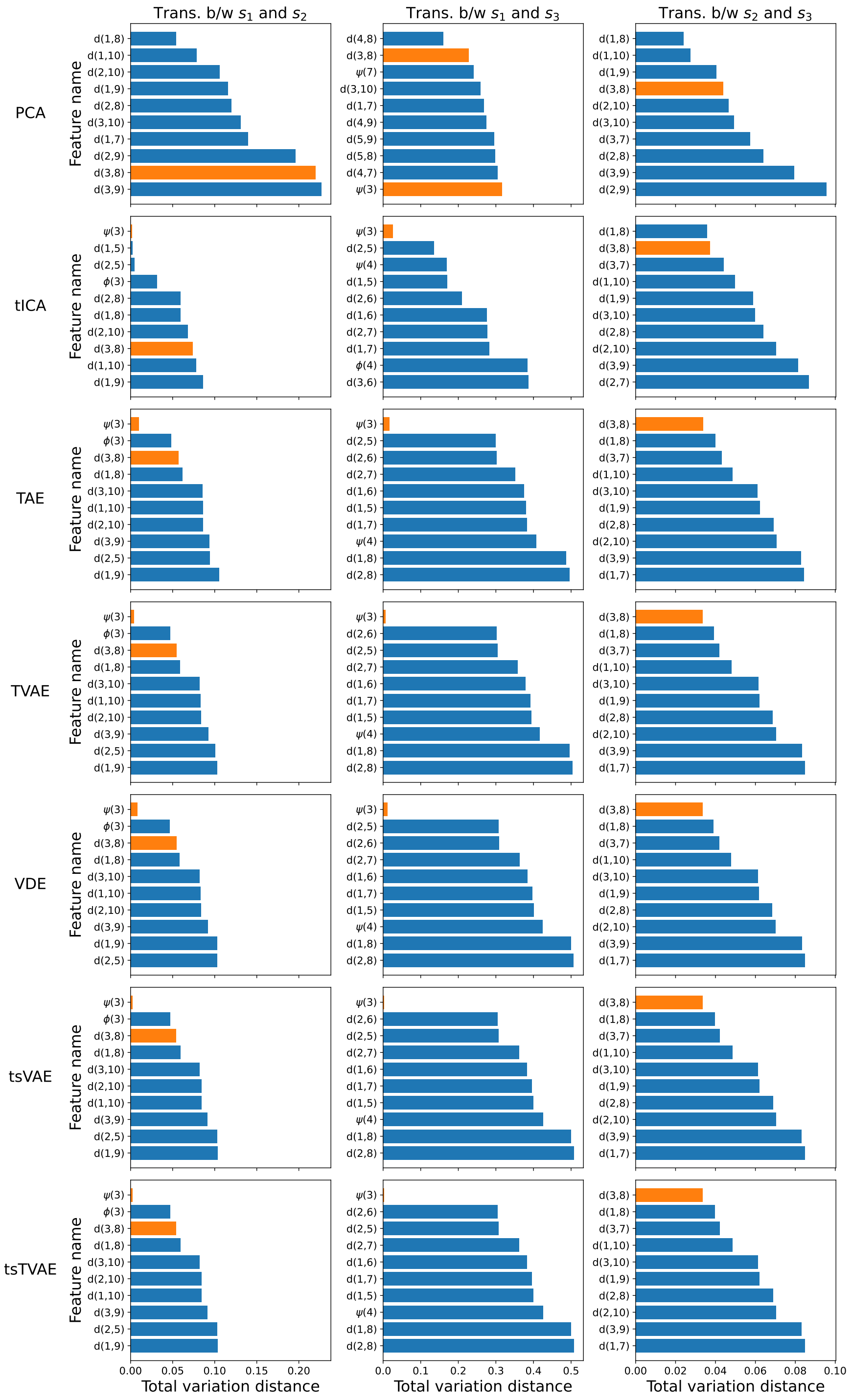


Figure 3: Total variation distance between the histograms of each variable of macro-states for chignolin trajectories. The variables include the distances between the no-adjacent residues and the dihedral angles of the residues. The ten smallest total variation distances for the transitions are displayed. The *orange* bar means the specific variables: the distance between ASP3N and THR8O and the dihedral angle $\psi$ of ASP3.

where $(\Omega, \mathcal{F})$ represents the measurable space. When $\Omega$ is a bin set of histograms, the total variation distance is computed by

$$\|p_1 - p_2\|_{\mathrm{TV}} = \sum_{i=0}^{N-1} |p_1(x + i\Delta x) - p_2(x + i\Delta x)|\Delta x, \tag{5}$$

where $\Delta x$ is the bin-size and $N$ is the the number of divisions.

Figure 3 shows the total variation distances between the histograms of each variable of macro-states for chignolin trajectories. Let $h_i(x)$ represents histograms of a variable $x$ of the $i$-th macro-state, i.e.,

$$h_i(x) = \text{histogram of } \{x_t | S_t = s_i,\ t \in \{1, \cdots, T\}\}, \tag{6}$$

where $S_t$ and $s_i$ represents the macro-state at a time $t$ and the $i$-th macro-state. We created the histograms whose bin edges are estimated by the Freedman Diaconis Estimator. We used the variables as the distance between the no-adjacent pairs of the residues and the dihedral angles of the residues.

This figure shows critical variables for each transition between macro-states. The total variation distance between the histogram of the distance $d(3, 8)$ of macro-states $s_2$ and $s_3$ is the lowest in TAE, TVAE, VDE, tsVAE, and tsTVAE. Since the distance is the critical variable for folding dynamics, these methods can explain the folding dynamics by the transition between $s_2$ and $s_3$. The total variation distances of $\psi(3)$ between $s_1$ and $s_3$ of the methods excluding PCA are the lowest. From these results, the NN-based methods can construct interpretable embedding space regarding the transition of macro-states.

### References


(1) Rose, C.; Murray, D. *Mathematical Statistics with MATHEMATICA*; Springer, 2002.

(2) Kruschke, J. *Doing Bayesian data analysis: A tutorial with R and BUGS*; Academic

Press / Elsevier, 2011.
